# Evolution of frost and drought responses in cool season grasses (Pooideae): was drought tolerance a precursor to frost tolerance?

**DOI:** 10.1101/2024.04.19.590374

**Authors:** Sylvia Pal Stolsmo, Camilla Lorange Lindberg, Rebekka Eriksen Ween, Laura Schat, Jill Christine Preston, Aelys Muriel Humphreys, Siri Fjellheim

## Abstract

Frost tolerance has evolved many times independently across flowering plants. However, conservation of several frost tolerance mechanisms among distant relatives suggests that apparently independent entries into freezing climates may have been facilitated by repeated modification of existing traits (‘precursor traits’). One possible precursor trait for freezing tolerance is drought tolerance, because palaeoclimatic data suggest plants were exposed to drought before frost and several studies have demonstrated shared physiological and genetic responses to drought and frost stress. Here, we combine ecophysiological experiments and comparative analyses to test the hypothesis that drought tolerance acted as a precursor to frost tolerance in cool-season grasses (Pooideae). Contrary to our predictions, we measured the highest levels of frost tolerance in species with the lowest ancestral drought tolerance, suggesting that the two stress responses evolved independently in different lineages. We further show that drought tolerance is more evolutionarily labile than frost tolerance. This could limit our ability to reconstruct the order in which drought and frost responses evolved relative to each other. Further research is needed to determine whether our results are unique to Pooideae or general for flowering plants.

**Highlight:** We tested whether drought tolerance was an evolutionary precursor to frost tolerance in grasses (Pooideae), but found these responses to be negatively correlated, suggesting they evolved independently in different lineages.

## Introduction

Two thirds of the global land area experiences frost at least some time during the year (Larcher, 2005). Frost is one of the most severe abiotic stresses plants can experience and the inability to cope with frost is thought to limit the distribution of many species (Donoghue, 2008). Based on this, it is widely held that evolutionary transitions from tropical, frost-free environments to those experiencing freezing are difficult. Despite this, temperate species are found in many angiosperm lineages (ca. 40% of families), and frost tolerance appears to have evolved multiple times independently (Preston and Sandve, 2013; Ricklefs and Renner, 1994; Schubert *et al*., 2020; Watcharamongkol, Christin and Osborne, 2018) (Kottek *et al*., 2006; Stevens, 2001). One caveat of this view is that some cold stress responses are conserved across distantly related species and similar ancestral pathways have repeatedly been involved in their evolution (Preston and Sandve, 2013; Schubert *et al*., 2019a). This suggests that the origin of frost tolerance in different lineages may not have been truly independent, but may instead have occurred by repeated modification of the same ancestral stress tolerance responses. Such ancient stress tolerance pathways may therefore have acted as precursors, or exaptations, to the sophisticated frost tolerance responses of many lineages today.

The most obvious candidate for an evolutionary precursor to frost tolerance is some form of drought tolerance. In general, strategies for avoiding dehydration are thought to be more ancient than adaptations to low temperature stress. All land plants need some basic mechanism for avoiding dehydration and some drought tolerance responses are ancient, most likely having their origins early during land plant terrestrialisation or the evolution of vascular plants, some 400–500 million years ago (Mya) (Bowles, Paps and Bechtold, 2021; Oliver, Tuba and Mishler, 2000; Preston and Sandve, 2013; Sakai and Larcher, 1987a). In contrast, while cool-climate pockets may have been present in Northern Hemisphere mid-latitude mountain areas in the Eocene (56–34 Ma; (Hagen *et al*., 2019), emergence of cold and freezing environments of today is not thought to have begun until the late Eocene (mainly from ca. 34 My; (Eldrett *et al*., 2009; Liu *et al*., 2009; Pound and Salzmann, 2017; Zachos *et al*., 2001a). Thus, flowering plants are thought to have evolved in a relatively warm world, with traits for dealing with frost stress evolving by independent repurposing of ancestral stress pathways (Preston and Sandve, 2013; Schubert *et al*., 2019a; Schubert *et al*., 2020).

The idea that there is a mechanistic link between adaptations to frost and drought was first put forward by Ebermayer (Ebermayer, 1873) in his ‘frost desiccation theory’. Ebermayer realized that both drought and frost stress require tolerance of cellular desiccation, and there is now ample evidence supporting the fact that the water deficit caused by both drought and freezing elicits many common physiological responses (Preston and Sandve, 2013; Sakai and Larcher, 1987b; Shinozaki and Yamaguchi-Shinozaki, 2000; Shinozaki, Yamaguchi-Shinozaki and Seki, 2003). For example, both drought and frost can cause cells to collapse. Under freezing conditions, this is caused by ice crystal formation, either intracellularly, leading to mechanical puncturing of cell membranes, or extracellularly, leading to water withdrawal from the cells and causing them to shrink and collapse (Pearce, 2001). During drought, water deficit is the result of little to no available water or moisture. When the protoplast shrinks as a consequence of this, the concentration of cellular solutes increases above normal levels and, when the desiccation has reached a certain point, the cell collapses (Larcher, 2005).

Resistance to low water content in the cells can be induced by the synthesis and accumulation of solutes (Monson *et al*., 2006; Streeter, Lohnes and Fioritto, 2001), as well as through fortification and waterproofing of cell walls to protect the cell membrane against physical damage. These processes cause the intracellular water content to decrease (Vicre, Farrant and Driouich, 2004), which increases the cells’ ability to maintain turgor at lower leaf water potential (Monson and Smith, 1982), leading to increased tolerance of both drought (Engelbrecht and Kursar, 2003) and frost (Anisko and Lindstrom, 1996). Furthermore, both accumulation of solutes and increased cell wall thickness raise the plant dry matter content; there is thus often a correlation between leaf dry matter content and resistance to desiccation (Cornelissen *et al*., 2003). Accordingly, a positive relationship between leaf dry matter content and both frost and drought tolerance has been reported in various plants from different environments (Pescador *et al*., 2016) (Liu *et al*., 2015). However, it is unclear whether high leaf dry matter content specifically confers tolerance of both drought and frost, or whether this is a general effect of greater desiccation resistance.

A physiological link between drought and frost stress has also been demonstrated in the field. Plants that have been exposed to drought and then subjected to frost show increased frost tolerance, whereas pre-treatment with heat had no effect on subsequent frost tolerance (Pisek and Larcher, 1954; Sumner *et al*., 2022). Similarly, acclimation to freezing can result in acclimation to drought, and vice versa (Hussain *et al*., 2018; Medeiros and Pockman, 2011), and plants from humid mountains often have lower frost tolerance than plants from arid mountains, even though the arid mountains are no colder than the humid ones (Sierra-Almeida, Reyes-Bahamonde and Cavieres, 2016). Together, these studies suggest that stress pathways activated during one type of stress can yield physiological responses that are beneficial during the other. However, there has been little research addressing how the positive relationship between drought and frost responses evolved (see also (Folk, Siniscalchi and Soltis, 2020; Folk *et al*., 2019).

Pooideae (Poaceae) are a globally distributed clade of cool season grasses dominating arctic, continental and temperate floras (Hartley, 1973; Visser *et al*., 2014; Zhang *et al*., 2022). These habitats are characterized by short growing seasons, high temperature and precipitation seasonality, as well as episodic (short term/diurnal) and periodic (seasonal) frost and drought events. Pooideae are also distributed in a range of other environments, including forests, deserts and saline areas, with some lineages (e.g. Meliceae) being found in warmer and moister habitats and others (e.g. Triticeae) in colder and drier habitats (Bennett, Flowers and Bromham, 2013; Kellogg, 2015; Zhang *et al*., 2022). The global distribution of Pooideae across such divergent habitats makes the group well suited for testing how adaptations to dry and freezing environments evolved relative to each other.

The ancestors of Pooideae most likely evolved as forest understory plants during a warm period in the late Cretaceous (77–58 Mya) (Bouchenak-Khelladi *et al*., 2010; Gallaher *et al*., 2019; Gallaher *et al*., 2022b; Kellogg, 2001; Schubert *et al*., 2019b; Zachos *et al*., 2001b). It is thought that Pooideae transitioned from closed forest environments to open habitats several times independently, but it is unclear exactly how many times and in which lineages. Previous studies suggest at least two independent transitions to open habitats, in the most recent common ancestors of the core Pooideae (Poeae + Triticeae) and Stipeae, or in the lineages leading up to these clades (Elliott *et al*., 2023; Zhang *et al*., 2022). Additional independent transitions may also have occurred in other lineages (e.g. in Nardeae and Lygeeae). These transitions began 67–58 Mya (Gallaher *et al*., 2022a; Schubert *et al*., 2019b; Strömberg, 2011), with the most likely drivers being increased aridity and altered disturbance regimes, e.g. seasonality, fire and herbivory, rather than global cooling (Edwards *et al*., 2010; Strömberg, 2011). Consequently, multiple lineages of Pooideae must have adapted to arid conditions early in their evolution. It has also been shown that sets of the same drought tolerance genes are expressed by distantly related species of Pooideae in response to short-term cold exposure (Schubert *et al*., 2019a). This supports the idea that ancient drought responses facilitated the evolution of early frost responses.

It is unclear to what extent early Pooideae were exposed to cold conditions. Even though the climate was generally warm, previous reconstructions suggest that Pooideae originated in the Palaeoarctic, in cool-climate pockets in the emerging Eurasian alpine orogeny. They were therefore possibly exposed to short-term cool conditions, but not prolonged winters (Das *et al*., 2023; Gallaher *et al*., 2022b; Schubert *et al*., 2019b). However, there remains considerable uncertainty regarding the biogeographical origins and the level of cold exposure experienced by early Pooideae. What is more certain is that from ca. 34 Mya onwards, Pooideae experienced progressively cooler and drier conditions while temperate biomes expanded (Pound and Salzmann, 2017; Zachos *et al*., 2001b). Accordingly, Pooideae is known to have successively evolved phenological and physiological adaptations to frost and short growing seasons (Fjellheim, Boden and Trevaskis, 2014; Fjellheim *et al*., 2022; McKeown *et al*., 2016; Sandve *et al*., 2011; Zhong *et al*., 2017), allowing them to diversify in northern temperate regions (Bouchenak-Khelladi *et al*., 2010; Kellogg, 2001; Schubert *et al*., 2020).

Here, we ask whether drought tolerance could have acted as an evolutionary precursor to frost tolerance during the evolution of Pooideae, by combining ecophysiological experiments with reconstructions of how drought and frost tolerance responses evolved using comparative, phylogenetic analyses. Specifically, we test the predictions that (1) drought and frost responses are positively correlated, (2) leaf dry matter content is positively correlated with drought and/or frost tolerance, (3) frost tolerant species are nested in ancestrally drought tolerant clades, (4) frost tolerance evolved more frequently in ancestrally drought tolerant than drought sensitive clades, and (5) climate conditions in species’ natural habitats can explain variation in drought and frost responses. Contrary to predictions, we find that frost and drought tolerance responses are negatively correlated, with frost tolerance being highest in ancestrally drought sensitive clades, high leaf dry matter content is associated with drought tolerance but not frost tolerance, while climate of origin is largelly unrelated to either.

## Materials and methods

### Species selection

Sixty two accessions representing 61 species were included in the experiment, sampled based on seed availability and their climatic, geographic and phylogenetic diversity. Two accessions of *Phleum pratense* were included, one identified simply as *P. pratense* and one identified as *P. pratense* ssp *nodosum*. The sampled species are mainly perennials and represent six of the ten tribes of Pooideae, including all of the major ones (Table 1) (Soreng *et al*., 2017) (Clayton *et al*., 2002 onwards). Species names follow accepted names according to The Plant List (2013).

**Table 1.**
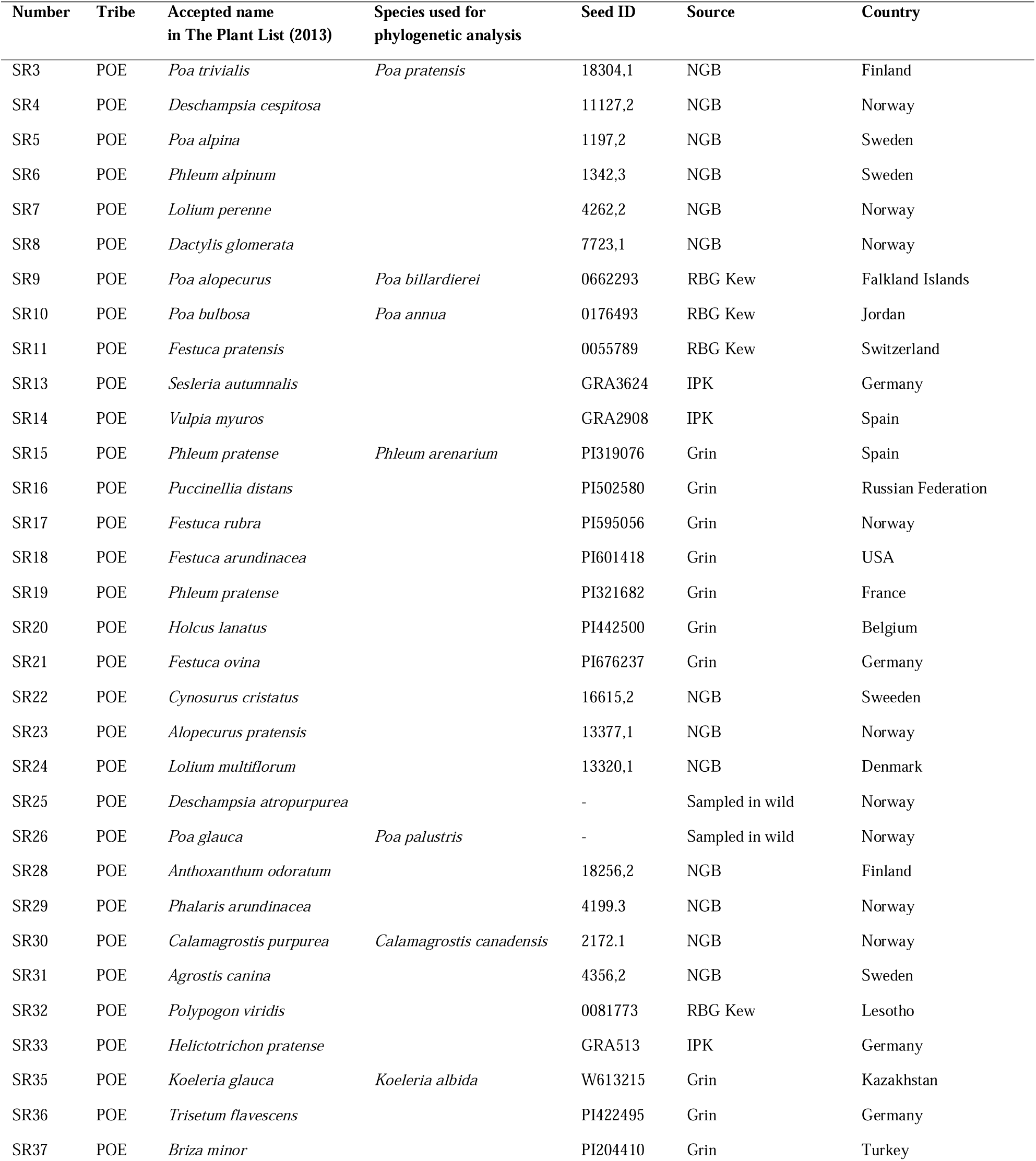
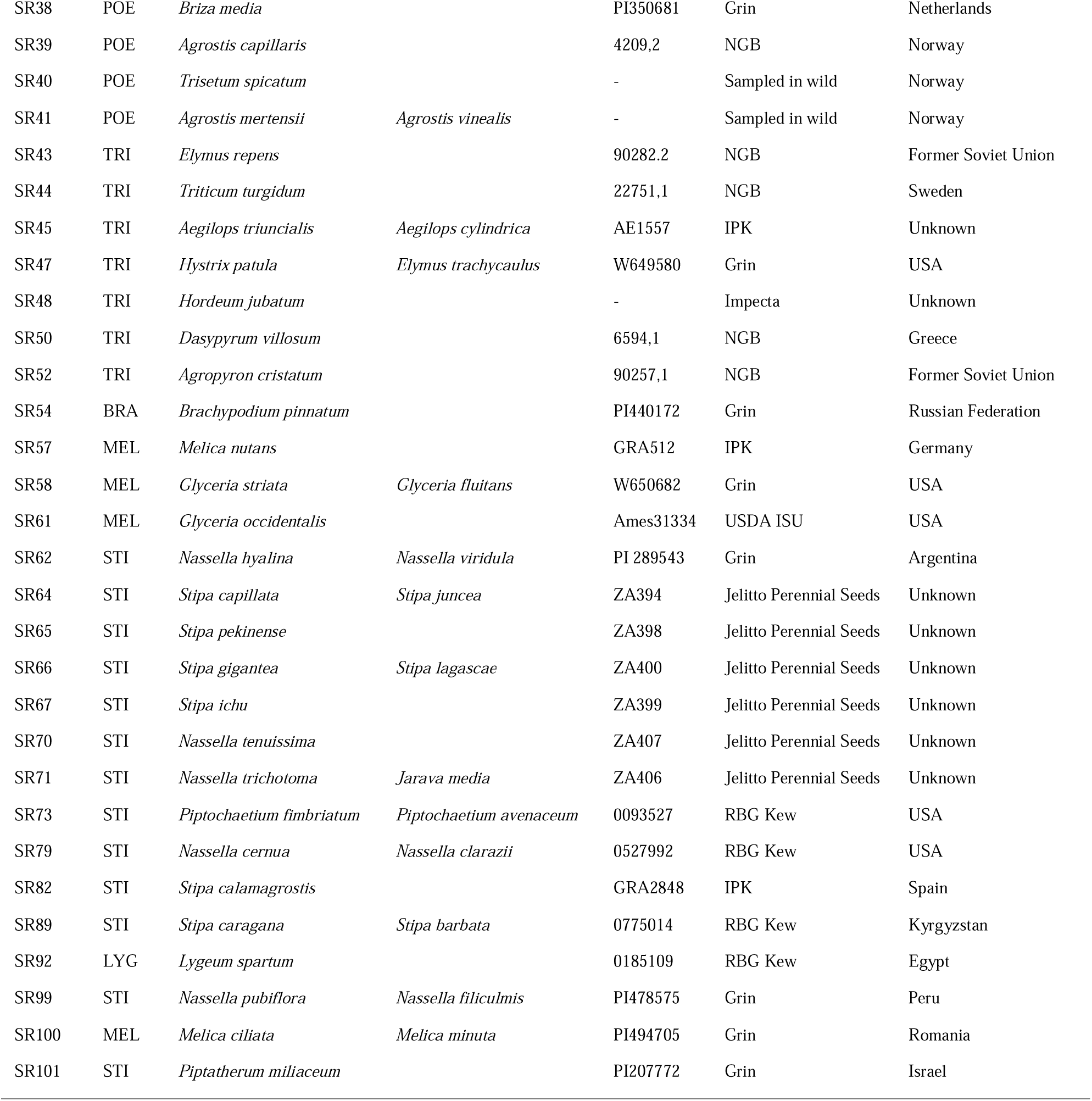
Species analysed in the experiment. The table shows the experimental population number, tribe, accepted scientific name, species from (Schubert *et al*., 2019b) used as phylogenetic placeholders in phylogenetic analyses, seed ID, source of the seeds and the country of origin. Tribes are abbreviated as: POE = Poeae, TRI = Triticeae, BRA = Brachypodieae, MEL = Meliceae, LYG = Lygeeae, and STI = Stipeae.

### Germination and growth

The experiment took place in a greenhouse at Vollebekk, Ås, Norway (59°39’42.4”N 10°45’01.5”E) from 14^th^ of September until 14^th^ of December 2018. The greenhouse held an average temperature of 17 °C and long day conditions with 16 hours of light. The light (200 mmol) was a mix of natural light through the windows and light from metal halide lamps with both Philips MASTER HPI-T Plus (400W/645 E40 1SL) and Osram POWERSTAR HQI-BT (400W/ D PRO) light bulbs.

To promote synchronized germination, seeds were stratified in humid soil at 4 °C for four days and then transferred to 25 °C for 24 hours, all in darkness. The seeds were then transferred to the greenhouse for germination. When plants were large enough (∼5 cm, approximately two to three weeks after germination), single tillers were pricked out in 8×8 cm^2^ pots filled with standard potting soil (“Gartner jord”, Tjerbo Torvfabrikk, Rakkestad, Norway). Forty-eight individuals per species were used in the experiment where possible (Table S1). Eighteen species had fewer than 48 individuals (n=29–47), which resulted in a total of 2,870 plants in the experiment (Table S1).

After pricking out, plants were watered once with fertilised water containing a mix of 800 g/100L YaraTera Kristalon Indigo (9 % N + 5 % P + 25 % K, Yara, Oslo, Norway) and 600 g/10L YaraLiva Calcinit (15.5 % N + 19 % Ca, Yara), in a solution with a conductivity of 1.7 mS/cm. Then, plants were grown for two weeks without fertilizer and then for one more week with daily watering with the fertiliser solution. Fertilisation was done to ensure that plants were robust at the start of the experiment and nutrients were not limiting regrowth after treatment. Plants were randomly rotated among the tables every week.

After the initial three-week growth period, plants were randomly divided into four treatment groups: (1) sudden frost at −1 °C, (2) sudden frost at −3 °C, (3) drought, and (4) control. Plants receiving sudden frost were moved directly from the 17 °C greenhouse to the frost conditions without acclimation. For most species, there were ten individuals in each treatment group, and four individuals each for initial electrolyte leakage and leaf dry matter content measurements (Table S1). Plants were randomly distributed in trays. Both the drought and frost treatments started on October 22^nd^ 2018.

### Drought treatment

The drought treatment took place in the greenhouse at Vollebekk with the light and temperature conditions described above. Since species have different rates of water uptake (Taiz *et al*., 2015) and the soil content might differ slightly between pots, soil moisture was measured in all pots during the drought treatment. The drought zone was defined as ≤ 5 % soil moisture. A HH2 Moisture Meter (Delta-T Devices Ltd, Cambridge, UK) with a Wet-2-sensor was used to measure soil moisture by placing it in the soil. To avoid taking measurements in the holes in the soil left from the previous measurement, which could influence the moisture reading, repeat measurements in the same pot were taken on opposite sides. In the case of large variation between the two measurements, a measure was taken at a third corner of the pot and the average was used.

Soil moisture was measured at the onset of the drought treatment and then every fourth day until the end of the treatment. To determine when the plants entered the drought zone, we the soil moisture decline rate, estimated using the initial and last soil moisture measurement of ≤ 20 % in formula (1):

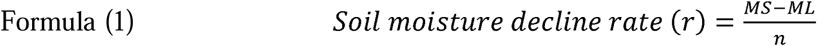

where *MS* is the starting moisture, *ML* is the last moisture recorded and *n* is number of days. The soil moisture decline rate was then used to estimate an approximate date when soil moisture was ≤ 5 % (formula (2).

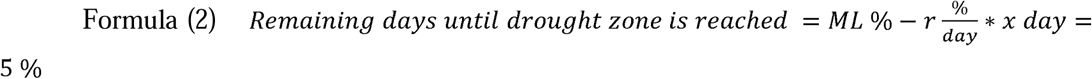

where *r* is the soil moisture decline rate found by using formula (1) and *x* is the number of days until the species hits the drought zone. Plants stayed in the drought zone for 4-5 days.

After the end of the drought treatment, leaves of 10 individuals per species were harvested for conductivity measurements (see below) and the plants were watered and cut down to approximately 2-4 cm height. Regrowth was scored after two and three weeks (see below).

### Sudden frost treatment

The sudden frost treatment took place in frost chambers (Weiss Umwelttechnik GMBH, model KWP 1000/55-10DU-S) at “Centre for Plant Research in Controlled Climate”, Ås, Norway (59°40’08.7”N 10°46’07.6”E) without additional light other than from the windows in the chambers. Minimum temperatures for mild and severe sudden frost were −1 °C and −3 °C, respectively. Following the protocol of Alm *et al*. (2011), the starting temperature was set to 0 °C for 12 hours and then lowered by 1 °C per hour to the minimum temperature, where it was kept for 24 hours. Then the temperature was increased by 1 °C per hour back to 0 °C. The plants remained at 0 °C for 24 (−1°C treatment) or 28 (−3 °C treatment) hours. Thereafter, the plants were watered and placed in a room at +3 °C to thaw. Leaves from four individuals per species were sampled and electrolyte leakage was measured (see below). After 24 hours at +3 °C, plants were moved back to the greenhouse and cut down to approximately 2-4 cm in height. Regrowth was scored after two and three weeks (see below).

### Control

Sudden frost and drought treatments were carried out simultaneously, which allowed for the use of the same control for both treatments. Control plants were kept in the greenhouse, under the conditions described above. Plants randomized on the tables and watered every week. Control plants were cut down to 2-4 cm, their regrowth was scored after two and three weeks, and their electrolyte leakage measured (see below).

### Ecophysiological measurements

#### Leaf dry matter content (LDMC)

To be able to correlate drought and frost tolerance responses with LDMC, four individuals per species were used to measure LDMC on the same day as the drought and frost treatment started. Fresh aboveground biomass was weighed for each plant, before being placed in individual paper bags, dried in a Unitherm drying oven (Russell-Lindsey Engineering Ltd., Birmingham, UK) at 90 °C for 14 hours and weighed again. LDMC was calculated as

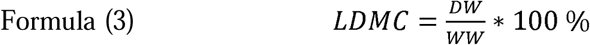

where *WW* is wet weight and *DW* is dry weight.

#### Electrolyte leakage and conductivity measurements

To assess the damage caused by the treatments, electrolyte leakage/conductivity was measured before and after the drought and frost treatments for all plants, including controls. When a cell gets damaged, it will release electrolytes (Hincha *et al*., 1987). Conductivity (mS) is a measure of amount of electrolytes released by a damaged leaf. High conductivity indicates high cell damage. Approximately 1 cm^2^ of a representative leaf was cut and placed in a tube with 10 mL distilled water. The samples were shaken at room temperature for ten hours before the conductivity was measured with a CWO Volmatic Mesur EC (Senmatic A/S DGT Volmatic, Søndersø, Denmark). The conductivity of the shaken samples was then divided by the maximum conductivity (formula (4)). To obtain the maximum electrolyte leakage per species for comparison, leaf samples were then boiled at approximately 97 °C for 11 minutes and the conductivity was measured again when the tubes had cooled down to room temperature (25 °C). To get percentage conductivity after each treatment, formula (4) was used per individual per species:

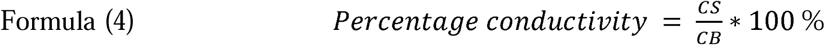

where *CS* is conductivity after shaking and *CB* is conductivity after boiling. To see if the treatments had any effect compared to the control group, formula (5) was used (Fujikawa and Miura, 1986):

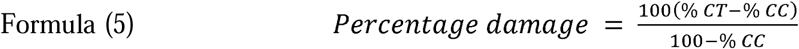

where *% CT* and *% CC* are the percentage conductivity obtained using formula (4) for the treatment and control groups respectively.

#### Fluorescence measurements

Fluorescence was measured on the control and drought plants before the start of the treatment and every fourth day until the plants had stayed in the drought zone for 4-5 days. The drought and control groups were measured on the same day for all species. Fluorescence measurements were carried out using FluorPen FP100 (Photon Systems Instruments, Drasov, Czech Republic) with the OJIP fluorescence transient analysis program. This program measures Fv/Fm, which represents the maximum quantum yield of photosynthetic efficiency in photosystem II. If the value of Fv/Fm is low, it can indicate that the plant is damaged due to low photosynthesis (Gilbert and Medina, 2016). The measurements were taken in the middle of a representative leaf per plant. To ensure an accurate measure of photosynthesis and to avoid light contamination, plants were placed in a dark room for 25-35 minutes before the fluorescence measurements were made in the dark. The plants were transferred to the dark three hours after dawn. Formula (6) was used to get the fluorescence of drought plants in relation to the control plants:

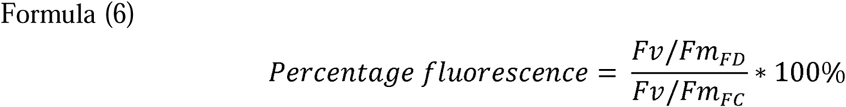

where Fv/Fm*_FD_*is the last measurement of the plant in the drought zone for each species before it was cut and Fv/Fm*_FC_* is the average measurements of the control for each species throughout the whole experiment.

#### Regrowth

Regrowth was assessed visually on a scale from 0 – 9, where 0 is dead and 9 is normal growth (Larsen, 1978). Formula (7) was used to obtain an estimate for the treatment plants in relation to the control plants:

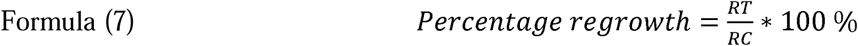

where *RT* is the average regrowth per species after two and three weeks for the treatment (RT = (RT_2weeks_+RT_3weeks_)/2) and *RC* is the average regrowth per species after two and three weeks for the control plants (RC = (RC_2weeks_+RC_3weeks_)/2).

### Statistical and phylogenetic analyses

#### Phylogenetic data

To analyse the experimental results in an evolutionary framework, information on the phylogenetic relationships among species was taken from Schubert et al. (Schubert *et al*., 2019b). The phylogenetic tree was pruned to retain only the species included in the experiment. Species in the experiment but not in the phylogeny (20 species) were assigned to the tips of their closest relatives (see Supplementary data 1). Phylogenies by Hamasha, von Hagen and Röser (2012) and Cialdella *et al*. (2007) were used to place species in tribe Stipeae, Grebenstein *et al*. (1998) was used for *Helictotrichon* and Gillespie, Archambault and Soreng (2007) for *Poa*. If no published phylogeny containing the species from both the experiment and its closest relative in the Schubert tree was found, the species in the experiment was assigned to a randomly selected tip within its respective genus (see Supplementary data 1). The species mean for each experimental variable (i.e. the seven measurements of drought and frost responses; Supplementary Data 1) was used to represent each species. All analyses were done with Rstudio version 1.1.383 (RStudio Team, 2016) and/or R version 3.5.2 (R Core Team, 2018).

#### Covariation and correlation among experimental variables

To visualise patterns of covariation among the seven experimental variables (Supplementary Data 1), we used a principal component analysis (PCA), performed using *ggbiplot* (Vu, 2011). Next, to test which experimental variables are statistically correlated with each other, pairwise regressions were performed using Pearson’s correlation test and the function ‘cor.test’. We also tested for autocorrelation among the residuals and, if detected, instead used a phylogenetic least-squares regression (PGLS), implemented with the ‘pgls’ function in *caper* (Orme *et al*., 2018). All pairwise trait combinations were tested.

#### Phylogenetic signal (λ)

To test whether closely related species showed more similar drought and frost responses than expected for a random sample of species, we estimated the phylogenetic signal of each trait. This information was used to select experimental variables for the evolutionary analyses below. Because Principal Component (PC) axes 2 and 3 showed interesting patterns suggesting covariation between frost and drought tolerance responses, we also tested whether there was phylogenetic signal in either of these variables. Phylogenetic signal was estimated by comparing the fit of different models with distinct assumptions for the variable Pagel’s λ (Pagel (1999). The Brownian Motion (BM) model assumes λ = 1, i.e. that the observed trait variance is completely correlated with the phylogenetic distance among species. The ‘white-noise’ model assumes λ= 0, i.e. that trait variance is independent of phylogeny. Finally, the ‘lambda’ model allows the value of λ to be estimated from the observed data, assuming a value between 0 and 1. The best model was determined based on the sample-size corrected Akaike Information Criterion (AICc (Akaike, 1974)), using a difference in AICc ≥ 2 to reject an inferior model (Anderson and Burnham, 2004). Models were fitted using *geiger* (Harmon *et al*., 2008).

#### Choosing experimental variables as proxies for drought and frost tolerance

We used the results of the pairwise correlations and estimates of phylogenetic signal to select experimental variables as proxies for drought and frost tolerance for further evolutionary analysis. We selected variables that were significantly correlated with other variables and that showed phylogenetic signal, because they convey information about several experimental responses and are more evolutionarily relevant. In this way “conductivity following drought treatment” was selected as a proxy for drought tolerance and “regrowth following the sudden frost treatment at −3 °C” as a proxy for frost tolerance. Leaf dry matter content (LDMC) was also analysed separately to test for a correlation with each proxy for drought and frost tolerance.

#### Ancestral state reconstruction

To visualise how drought and frost responses have evolved in Pooideae, ancestral states for the two proxy variables plus LDMC were reconstructed using the level of phylogenetic signal for each trait found above. This was achieved with the BM model, having first rescaled the phylogenetic branch lengths according to the phylogenetic signal of the trait in question (Table 2). Ancestral states were reconstructed under maximum likelihood (ML), using the ‘ace’ function in *ape* (Paradis and Schliep, 2018), and branches were rescaled using ‘rescale’ in *geiger* (Harmon *et al*., 2008). Finally, the reconstructed ancestral states were visualized using *ggtree* (Yu *et al*., 2017), *cowplot* (Wilke, 2019) and *ggplot2* (Wickham, 2016). Evidence for the expected patterns of drought tolerance evolving first and frost tolerance originating within ancestrally drought tolerant clades was assessed by eye.

**Table 2:** AICc values for the Brownian Motion (BM), ‘white’ (phylogenetically independent) and ‘lambda’ models and the value of lambda (λ) inferred under the best fitting model(s). The best fitting model(s) for each trait, with lowest AICc, is shown in bold. Models are considered indistinguishable if the difference in AICc < 2.

| | | AICc | | $\lambda$ |
| --- | --- | --- | --- | --- |
| MODEL/<br>VARIABLE | BM | WHITE | LAMBDA | (best<br>estimate) |
| SUDDEN FROST -1 °C |  |  |  |  |
| Regrowth | -28 | -51 | -61 | 0.47 |
| Conductivity | 513 | 447 | 449 | 0 |
| SUDDEN FROST -3 °C |  |  |  |  |
| Regrowth | 31 | 19 | 5 | 0.47 |
| Conductivity | 561 | 551 | 550 | 0 / 0.63 |
| DROUGHT |  |  |  |  |
| Regrowth | -50 | -64 | -62 | 0 |
| Conductivity | 640 | 602 | 603 | 0 / 0.11 |
| Fluorescence | 83 | 38 | 40 | 0 |
| PRINCIPAL<br>COMPONENT |  |  |  |  |
| PC2 | 240 | 211 | 213 | 0 |
| PC3 | 241 | 193 | 195 | 0 |
| LEAF DRY MATTER | 343 | 327 | 318 | 0.45 |

Relating species drought and frost responses and leaf dry matter content to local climate conditions

Finally, we tested whether species’ drought and frost responses and LDMC correlate with the climatic conditions in each species’ native environment. Because drought and frost responses may not only reflect adaptation to the local climatic conditions but can also bear signatures of species’ evolutionary and biogeographical histories (Coelho *et al*., 2019; Freckleton and Jetz, 2009; Humphreys and Linder, 2013; Lancaster and Humphreys, 2020), we included information on species’ phylogenetic relatedness and spatial proximity in these tests.

We used generalized least squares models, in which the variance in a response variable is partitioned into phylogenetic, spatial and independent components (denoted λ′, φ and γ, in turn) (Freckleton and Jetz, 2009). The phylogenetic (λ) and spatial (φ) components are estimated during model fitting. The relative contribution of phylogeny, λ′, is calculated from ML estimates of λ and φ, as (1 – φ) *x* (1 – λ). A high value (approaching 1) of either λ′ or φ indicates a large effect of phylogenetic relatedness or geographical proximity, respectively. If the relative contributions of spatial and phylogenetic distances do not sum to 1, then the remainder of the interspecific variation is independent of either geographical proximity or phylogenetic relatedness and can be related to other explanatory variables, such as local climatic variation.

We implemented a series of mixed effect linear regressions with phylogenetic and/or spatial distances as random effects and climate predictors as fixed effects. We defined drought tolerance (“conductivity following drought treatment”), frost tolerance (“regrowth following frost treatment at −3 °C”) and LDMC as response variables. As predictors we used four temperature (bio1, bio4, bio5 and bio6) and four precipitation (bio12, bio13, bio14 and bio15) variables from WorldClim2 (Fick and Hijmans, 2017), based on geographical occurrence data obtained from the Global Biodiversity Information Facility (GBIF). The eight BioClim variables were chosen to represent (average) annual conditions (bio1 – average annual temperature; bio12 – annual precipitation), upper and lower extremes (bio5 – maximum temperature of the warmest month, bio13 – precipitation of the wettest month, bio6 – minimum temperature of the coldest month, bio14 – precipitation of the driest month), and annual variation (bio4 – temperature seasonality, bio15 – precipitation seasonality). Geographical data were compiled by Schat et al. (unpublished; (GBIF.org, 2022a) see Supplementary Data S1), supplemented with data downloaded directly from GBIF for three species (*Achnatherum calamagrostis*, (GBIF.org, 2022b); *Lolium arundinaceum*, (GBIF.org, 2022c); and *Lolium pratense*, (GBIF.org, 2022d). GBIF occurrence records were filtered following Schat et al. (unpublished). From these we extracted the median latitude and longitude across each species range to represent the geographical centroids and median values for the BioClim variables to represent the climatic conditions in each species’ native range.

A spatial distance matrix was calculated using the ‘earth.dist’ function in the R package *fossil* (Vavrek, 2011) and a phylogenetic variance-covariance matrix was calculated using ‘vcv.phylo’ in *ape*. Moran’s *I* was also calculated using ‘Moran.I’ in *ape* to separately assess any spatial patterns in the data. Moran’s *I* is a correlation coefficient that ranges from −1 to 1, where 1 denotes perfect clustering of similar values, 0 is no autocorrelation (perfect randomness) and −1 is perfect clustering of *dissimilar* values (akin to perfect dispersion).

First, we fitted normal univariate linear regressions for each predictor and response variable to assess each climate variable’s effect on each predictor using the ‘lm’ function in R. Next, we fitted univariate mixed effect linear models, with phylogenetic and spatial distances as random effects and each climate variable as a fixed effect, partitioning the variance among phylogenetic, spatial and independent components. Finally, we proceeded with multivariate mixed effect models including just the predictors with the strongest effects in the univariate tests and calculating the variance partitioning into random effects as before. Mixed effect models were fitted using the ‘regress’ function in the R package *regress* (Clifford and McCullagh, 2006, 2014) and code from (Cardillo and Skeels, 2016). For the multivariate models we used AIC scores to compare the fit of a full model (including the predictors and spatial and phylogenetic distances) with the fit of a series of reduced models (including any combination of phylogenetic distance, spatial distance and predictors). We note that this approach does not allow estimation of how much of the *total* trait variance that can be attributed to phylogenetic and spatial distances relative to the local climate (equivalent to the R^2^ for a linear regression; see (Ives, 2018) and *cf.* (Lancaster and Humphreys, 2020)), but it does allow quantification of the importance of phylogeny and geography for explaining drought and frost responses, as well as assessment of whether there is an effect of the local climate when any phylogenetic and spatial effects are accounted for.

## Results

### Variation in drought and frost tolerance responses

Almost all individuals showed full regrowth after sudden frost at −1 °C and the drought treatment, whereas there was much more variation in regrowth after sudden frost at −3 °C, including no regrowth at all (Supplementary Data, S2). Conductivity following drought and sudden frost at −3 °C also showed a range of values, whereas most plants had low conductivity following frost at −1 °C. The maximum quantum yield after drought measured between 0-0.95, with most plants having intermediate fluorescence (mean=0.55, SD=0.29).

### Principal component analysis

The PCA showed covariation among several of the experimental variables (Fig. 1). The first four PCs explained 86 % of the variance. Overall, there were clear patterns of covariation among the different drought response measures and among the different frost response measures, but mixed patterns regarding how the drought and frost responses covaried with each other (they do for PC2, partly for PC3 but not for PC1). PC1 explained 33% of the variance and primarily depicted the expected pattern of covariation between conductivity and fluorescence following drought treatment (Fig. 1a). In addition, PC1 showed that drought tolerant species (low conductivity, high fluorescence) are less frost tolerant (low regrowth and high conductivity following frost treatment). PC2 (24% of the variance) had similar loadings from all experimental variables and showed that the three conductivity measures increase with increasing PC2, while all other measures decrease. Thus, species at the lower extreme of PC2 had high tolerance of both frost and drought. All the traits covaried in the same direction with the third PC (18 % of the variance; albeit with very low loadings for fluorescence and conductivity following drought treatment; Fig. 1b). Thus, lower extremes of PC3 mainly group frost tolerant species that are somewhat drought tolerant as well. The fourth PC (11 % of the variance; Fig. 1b) mainly covaried with regrowth after drought.

**Figure 1.**
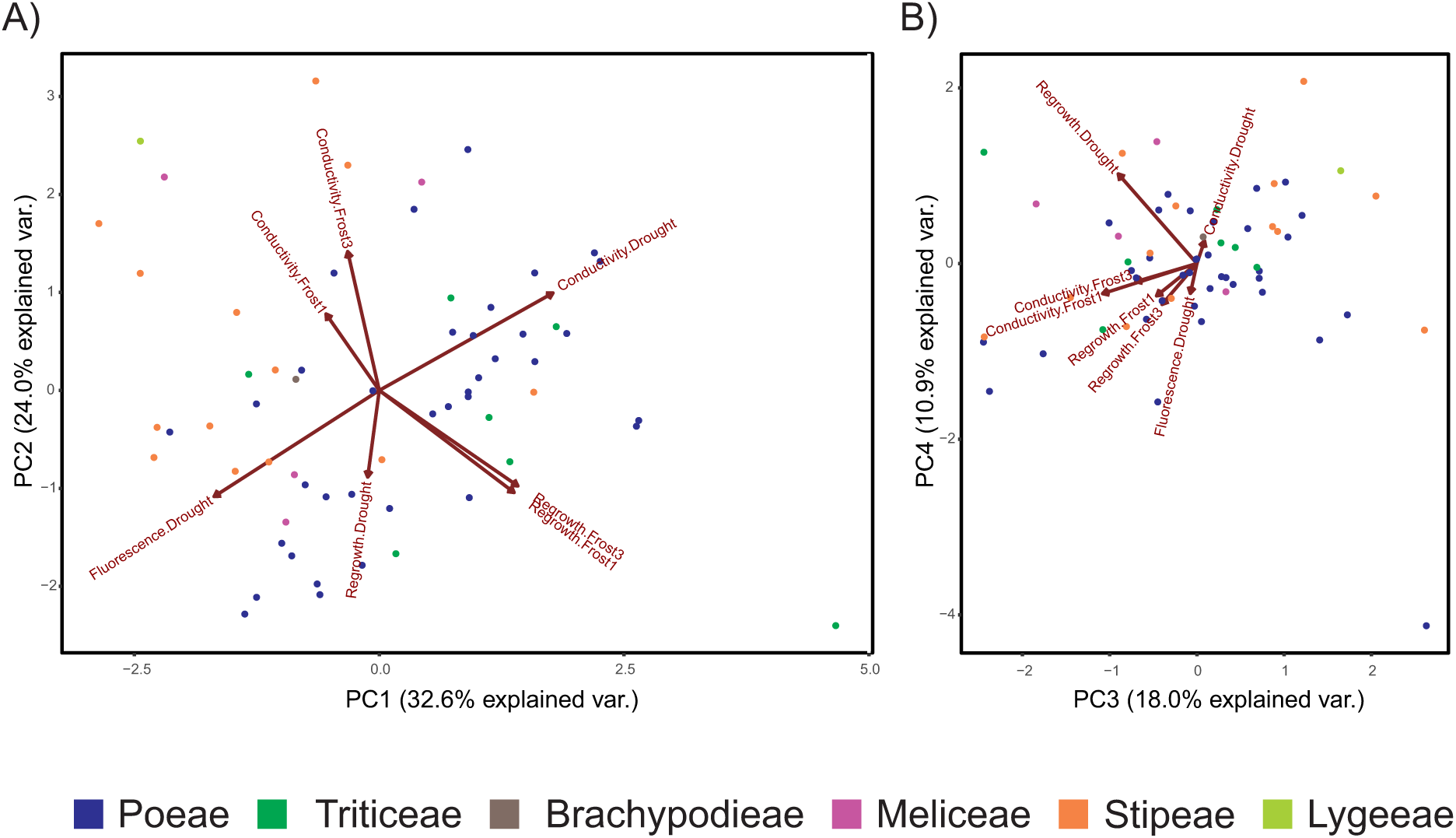
Principal component analysis (PCA) of the experimental variables. These plots are visualisations of the patterns of covariation in the data, as a means of data exploration. A) The first two principal components (PC1 and PC2), which together explain 57% of the variance. B) The third and fourth principal components (PC3 and PC4), which together explain 29% of the variance. Each dot represents an accession/species (n=62), coloured according to the tribe in which it is classified. The arrows labelled with the experimental variables show in which direction and by how much (length of the arrow) each variable contributes to the distribution of the species, in relation to the other traits. The direction of each arrow in relation to each PC axis also shows which variables contribute most strongly to each PC. The seven experimental variables included are: regrowth, fluorescence and conductivity following drought treatment and regrowth and conductivity following frost treatment at −1 and −3 °C (Supplementary Data 1). Overall, these plots show mixed patterns regarding how the drought and frost response measures covary with each other but species in the bottom left part of A) show high tolerance of both drought and frost.

### Pairwise correlation tests

Five pairwise correlation tests were significant: regrowth after sudden frost at −1 °C and −3 °C (PGLS, *P* << 0.001, R^2^ = 0.32); conductivity after sudden frost at −1 °C and −3 °C (*P* < 0.05, Pearson’s *r* = 0.39); regrowth following sudden frost (−3 °C) and conductivity following drought treatment (*P* < 0.05, Pearson’s *r* = 0.27); conductivity and fluorescence following drought treatment (*P* << 0.001, Pearson’s *r* = −0.90); and LDMC and conductivity following drought treatment (PGLS, P < 0.05, R^2^ = 0.16). Two further tests were marginally significant: regrowth following −1 °C and drought treatment (P=0.062) and LDMC and regrowth following −3 °C (P=0.072); we do not consider these tests any further. Thus, the only test suggesting a significant correlation between drought and frost responses indicates decreasing drought tolerance (increasing conductivity) with increasing frost tolerance (increasing regrowth). LDMC increased with increasing drought tolerance (decreasing conductivity) but showed no relationship with frost tolerance.

### Phylogenetic signal

The strongest phylogenetic signal was found for regrowth following frost treatment at both −1 and −3 °C (λ = 0.47 in both cases) and LDMC (λ = 0.45; Table 2). Some evidence for phylogenetic signal was also found for conductivity following frost (−3 °C, λ = 0.63) and drought (λ = 0.11) treatment but the lambda and white models were statistically indistinguishable for these variables (Table 2). No other variable showed phylogenetic signal.

### Ancestral state reconstruction

Conductivity following drought treatment and regrowth following frost treatment at −3 °C were used as proxies for drought and frost tolerance, respectively. Frost treatment at −3 °C distinguished the species responses better than treatment at −1 °C, resulting in a response measure with greater variance. The ancestral state reconstruction for drought tolerance showed that tribes Stipeae and Lygeeae were ancestrally more drought tolerant (lower conductivity; yellower internal nodes; Fig. 2), compared to the rest of the Pooideae (Meliceae, Brachypodieae, Triticeae and Poeae; greener internal nodes; Fig. 2). Stipeae and Lygeeae were also inferred to have lower ancestral frost tolerance (less regrowth; blue internal nodes; Fig. 2) compared to the rest of Pooideae (green internal nodes; Fig. 2), with slightly higher ancestral frost tolerance in Triticeae and the Poeae chloroplast 2 clade (sensu (Soreng *et al*., 2017; Soreng *et al*., 2015) relative to other clades (yellower-green ancestral shades; Fig. 2). Finally, Stipeae and Lygeeae were inferred to have higher ancestral LDMC compared to core Pooideae (Poeae and Triticeae; Fig. 3).

**Figure 2:**
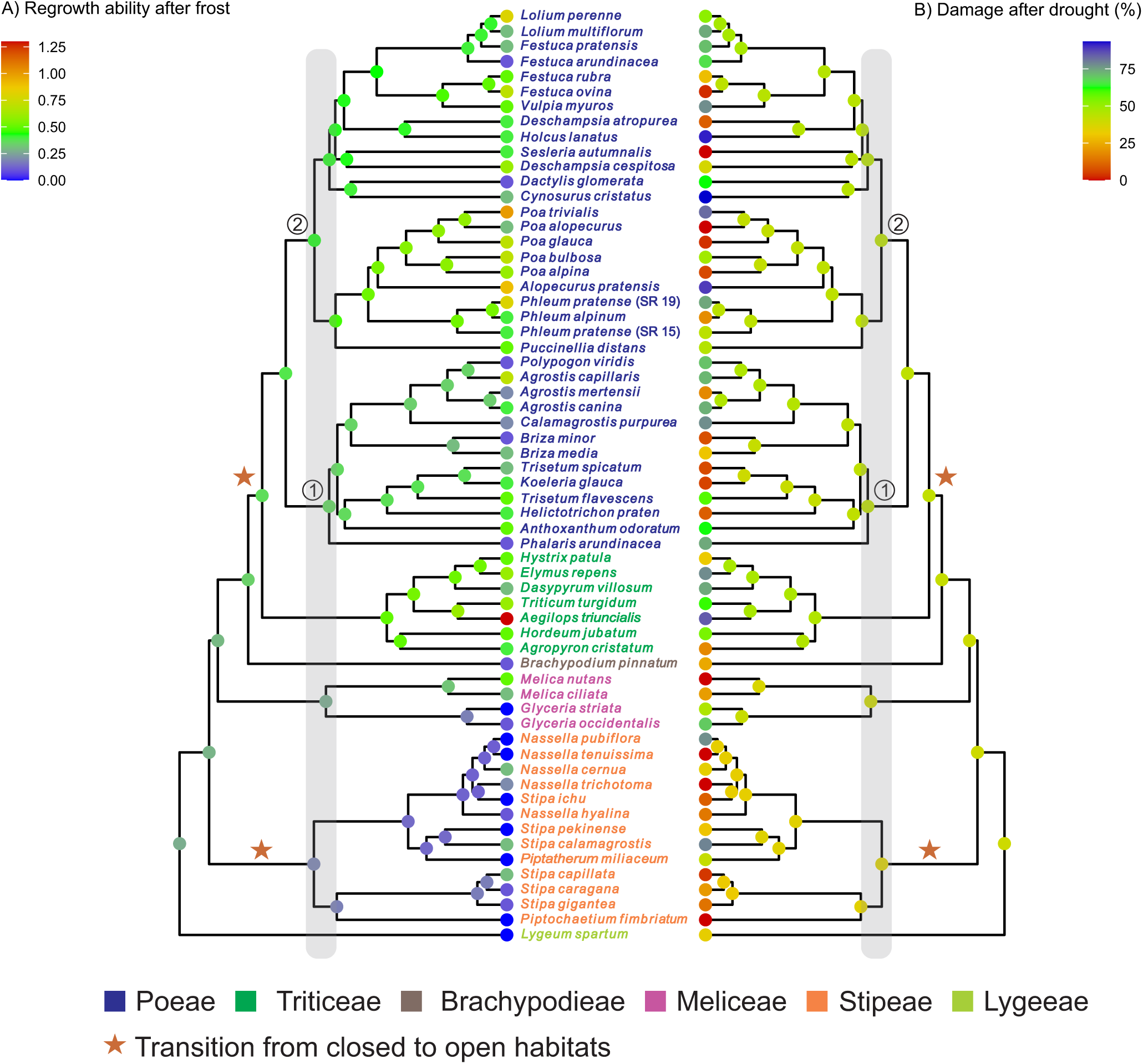
Ancestral state reconstructions of measured responses to frost and drought treatment based on a phylogenetic tree including all accessions/species in the experiment (n=62). A) Regrowth ability following frost treatment at −3 °C. B) Leaf damage (conductivity) following drought treatment. Values are expressed relative to the control. Node colours: reds indicate high levels of regrowth and low levels of damage/conductivity (i.e. high tolerance of frost and drought, respectively); blues indicate high levels of damage/conductivity and low levels of regrowth (i.e. poor tolerance of frost and drought, respectively). Overall, the ancestral state reconstructions show that high levels of frost tolerance (warmer colours, in A) evolved in clades that were ancestrally more drought sensitive (cooler colours, in B). Species names are coloured according to the tribe in which they are classified. Circled numbers indicate clades (“chloroplast subgroups”) as defined by Soreng *et al*. (Soreng *et al*., 2017). Grey shading indicates approximately the Eocene-Oligocene boundary at 34 Mya (molecular dates from (Schubert *et al*., 2019b)). Stars indicate putative transitions from closed to open habitats (Elliott *et al*., 2023; Zhang *et al*., 2022).

**Figure 3.**
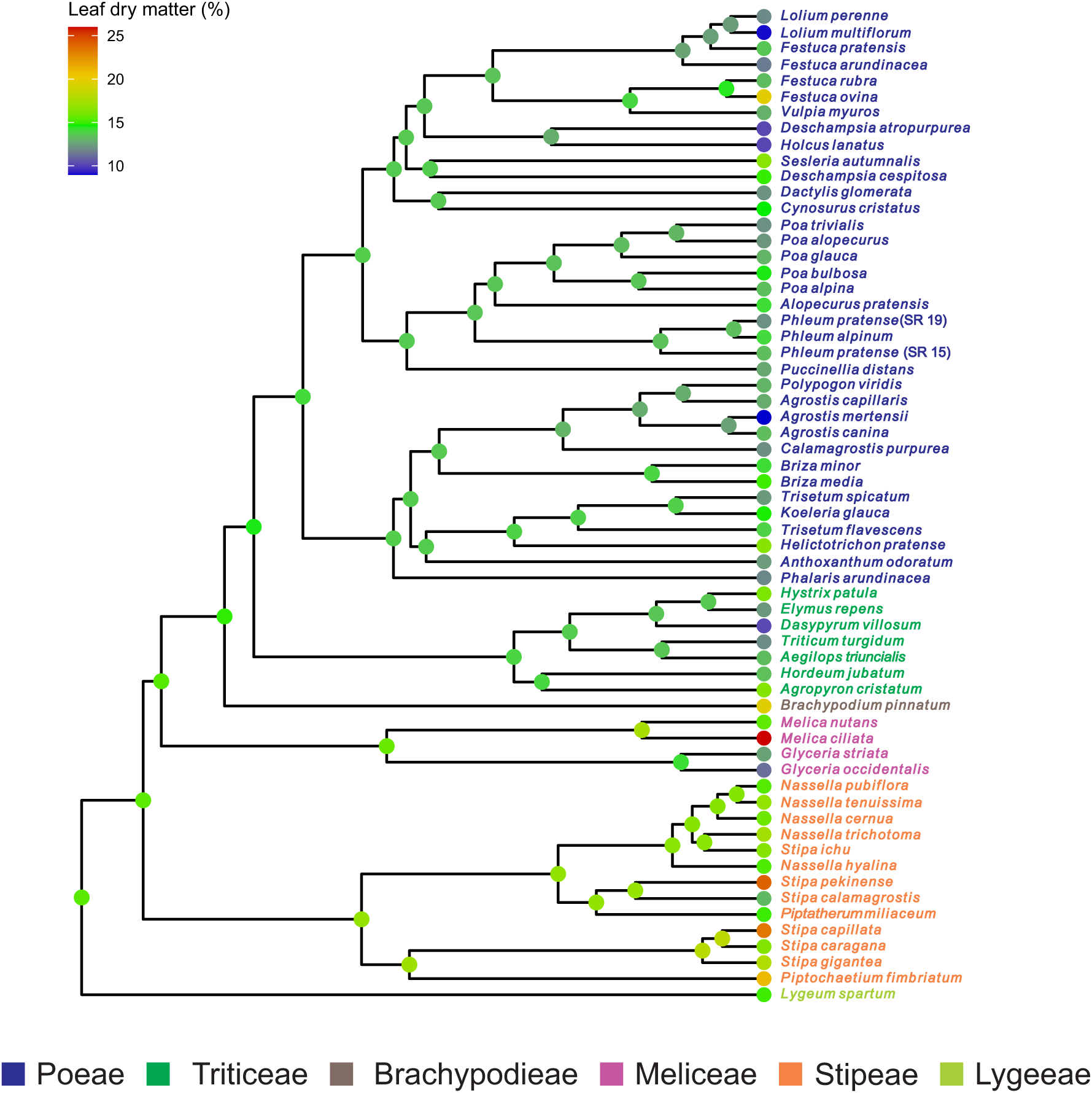
Ancestral state reconstruction for leaf dry matter content (LDMC) based on a phylogenetic tree including all accessions/species in the experiment (n=62). Node colours: reds indicate high LDMC; blues indicate low LDMC. Overall, the ancestral state reconstruction shows that the highest LDMC is found in clades that are the most drought tolerant, not frost tolerant (*cf.* Fig. 2). Species names are coloured according to the tribe in which they are classified.

### Spatial, phylogenetic and climatic correlates of drought tolerance, frost tolerance and leaf dry matter content

There was no evidence for a significant spatial signal in the analysed data. All values of Moran’s *I* were below zero, i.e. showing some degree of dispersion. This was significant for frost and drought tolerance (Moran’s *I* = −0.12 ± 0.0133, *P* < 0.001 [frost]; −0.075 ± 0.0132, *P* < 0.001 [drought]), but not LDMC (i.e. no spatial autocorrelation; Moran’s *I* = −0.029 ± 0.0133, *P* = 0.33).

The univariate linear regressions suggested that average climate conditions across each species range are poor predictors of how species responded to the drought and frost treatments. For frost tolerance, one climate variable had a significant effect (Bio15, precipitation seasonality; P =0.032, R^2^=0.075), whereas for drought tolerance and LDMC none was significant (P>0.05, results not shown). No adjustments for multiple testing were made.

The univariate mixed effect models with the variance partitioned into spatial, phylogenetic and independent components revealed that phylogeny explained almost all of the variance for all three response variables (λ′ > 0.999; Table 3). For frost tolerance, the strongest predictor effects were for three temperature variables (bio1, bio5 and bio6) but no test remained significant after correction for multiple testing (Table 3a). For drought tolerance, the strongest predictor effects were for three temperature variables (bio1, bio4 and bio6) and one precipitation variable (bio15), with all but bio1 remaining significant after correction for multiple testing (Table 3b). Finally, for LDMC, the strongest predictor effects were for one temperature (bio4) and two precipitation parameters (bio14 and bio15) but none remained significant after adjustment for multiple testing (Table 3c).

**Table 3.**
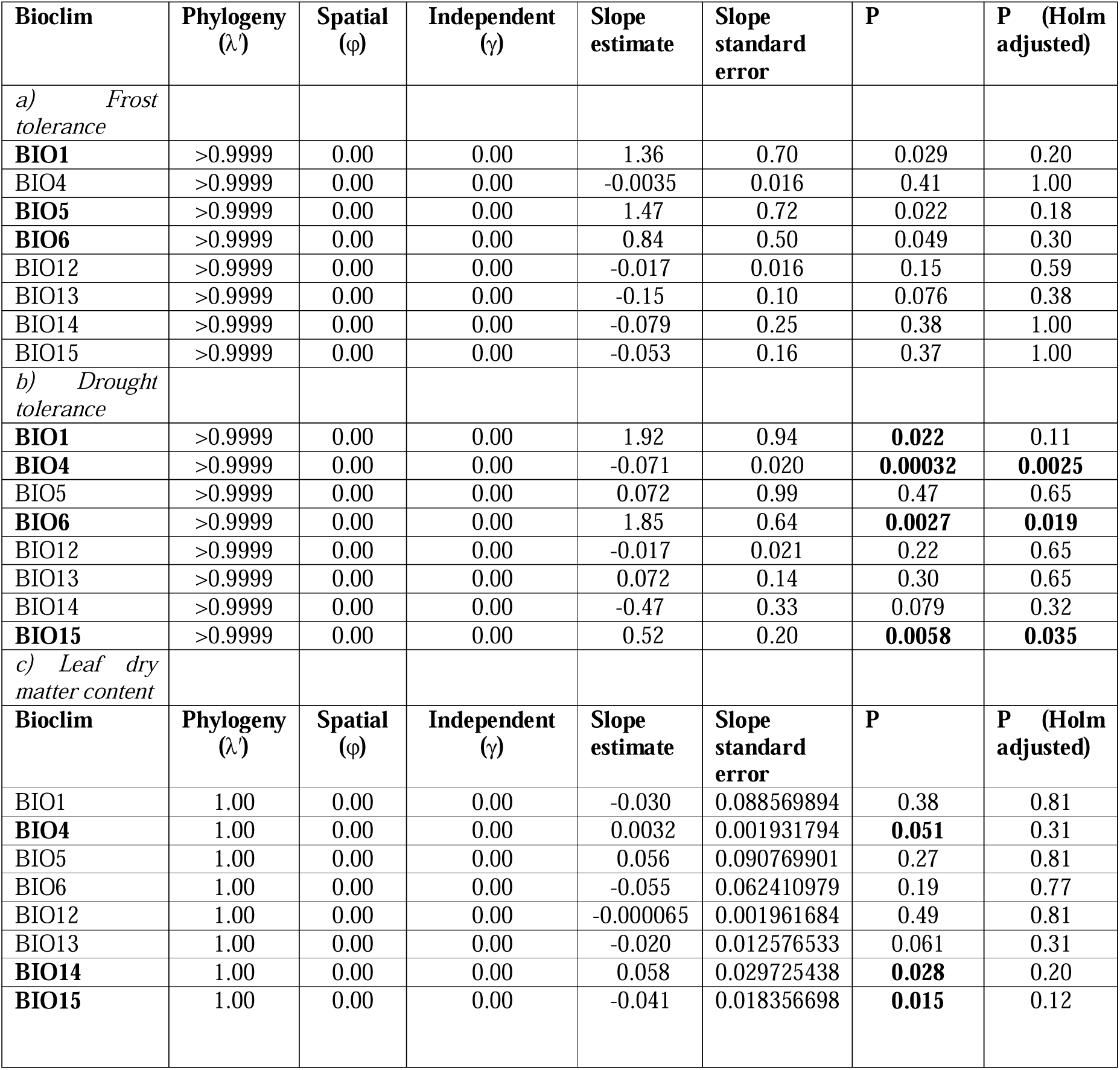
**Univariate mixed effect linear models with the variance partitioned into spatial (φ), phylogenetic (λ′) and independent (γ) components** and testing the effect of each BioClim predictor variable separately for a) frost tolerance (regrowth following −3 °C frost treatment), b) drought tolerance (conductivity following drought treatment) and c) leaf dry matter content. Significant tests are shown in bold.

The best multivariate mixed effect model for frost tolerance included just the predictors and the phylogeny (ΔAIC ≥ 12.6 compared to all other models; Table 4). Under this model, λ′= 0.50 and γ = 0.50, meaning that half the variance is attributed to phylogenetic distance and half is independent of either phylogenetic or spatial distance. However, none of the predictors, bio1, bio5 and bio6, showed a significant effect (P = 0.09, 0.08, 0.08, respectively; not shown) but removing the predictors from the model completely led to a much worse model (ΔAIC = 15.6; or ΔAIC = 12.4 for the full model vs. the spatial+phylogeny model; Table 4).

**Table 4.** Multivariate mixed effect linear models testing the effect of species’ local environment on measured frost and drought responses and leaf dry matter content (LDMC). Variance in the models is partitioned into some combination of spatial (φ), phylogenetic (λ′) and independent (γ) components. Predictors refers to several bioclimatic variables (see Methods). a) Frost tolerance (regrowth following −3 °C frost treatment). b) Drought tolerance (conductivity following drought treatment). c) LDMC. Best-fitting model(s) are shown in bold.

| | LL | AIC | ΔAIC | $\phi$<br>(spatial) | $\lambda'$<br>(phylogeny) | $\gamma$<br>(independent) |
| --- | --- | --- | --- | --- | --- | --- |
| *a) Frost tolerance* |  |  |  |  |  |  |
| Predictor + phylogeny + spatial | -225.7 | 465.3 | 27.0 | 0.00 | >0.9999 | <0.00001 |
| Predictor + spatial | -220.5 | 452.9 | 14.6 | 0.00 | – | 1.00 |
| **Predictor + phylogeny** | -213.2 | **438.3** | **0.00** | – | 0.50 | 0.50 |
| Spatial + phylogeny | -237.6 | 483.2 | 44.9 | 0.00 | >0.9999 | 0.0000006 |
| Phylogeny only | -223.9 | 453.9 | 15.6 | – | 0.50 | 0.50 |
| Spatial only | -233.0 | 472.0 | 33.7 | 0.00 | – | 1.00 |
| Predictor only | -220.5 | 450.9 | 12.6 | – | – | – |
| *b) Drought tolerance* |  |  |  |  |  |  |
| Predictor + phylogeny + spatial | -233.9 | 483.9 | 26.6 | 0.00 | >0.9999 | 0.0000014 |
| **Predictor + spatial** | -222.6 | **459.3** | **2.00** | 0.00 | – | 1.00 |
| **Predictor + phylogeny** | -221.8 | **457.6** | **0.30** | – | 0.30 | 0.70 |
| Spatial + phylogeny | -238.7 | 485.4 | 28.1 | 0.00 | 0.0000028 | >0.9999 |
| Phylogeny only | -237.8 | 481.6 | 24.3 | – | 0.19 | 0.81 |
| Spatial only | -238.7 | 483.4 | 26.1 | 0.00 | – | 1.00 |
| **Predictor only** | -222.6 | **457.3** | **0.00** | – | – | – |
| *c) Leaf dry matter content* |  |  |  |  |  |  |
| Predictor + phylogeny + spatial | 103.5 | 221.1 | 25.3 | 0.00 | 1.00 | 0.00 |
| Predictor + spatial | -99.0 | 210.0 | 14.2 | 0.00 | – | 1.00 |
| **Predictor + phylogeny** | -91.9 | **195.8** | **0.00** | – | 0.56 | 0.44 |
| Spatial + phylogeny | -109.7 | 227.4 | 31.6 | 0.00 | >0.9999 | <0.0001 |
| Phylogeny only | -97.0 | 200.0 | 4.2 | – | 0.49 | 0.51 |
| Spatial only | -103.7 | 213.0 | 17.2 | <0.0001 | – | >0.9999 |
| Predictor only | -99.0 | 208.0 | 12.2 | – | – | – |

For drought tolerance, three models were statistically indistinguishable from each other (0.30 < ΔAIC < 2.00; Table 4), the model including the predictors and just spatial distance, the model including the predictors and phylogenetic distance, and the model including only the predictors. However, under the spatial model, γ = 1.00 (i.e. all variance is independent of spatial distance) and under the phylogenetic model, λ′= 0.30 and γ = 0.70 (i.e. most variance is independent of phylogeny). Accordingly, the best model overall (albeit not significantly so) is the model including only the predictors (Table 4). Under this model, there is a significant effect of bio1 (P=0.035, slope=-6.64±3.59, t=-1.85) and bio6 (P=0.024, slope=7.36±3.65, t=2.02).

For LDMC, the best model was the one including the predictors and just the phylogeny (ΔAIC ≥ 4.2, Table 4). Under this model, λ′ = 0.56 and γ = 0.44, meaning that just over half the variance is attributed to phylogenetic distance and the rest is independent of either phylogenetic or spatial distance. None of the predictors showed a significant effect (P > 0.05, not shown). Accordingly, removing the predictors resulted in only a slightly worse model (ΔAIC = 4.2).

## Discussion

### Present-day drought responses are negatively correlated with responses to episodic frost

In keeping with our predictions, we found that responses to drought and frost are correlated in Pooideae. However, in contrast to our predictions, the nature of this correlation shows that the species most tolerant of episodic (short-term) frost were the least tolerant of drought. This is evident from the pairwise correlations among the experimental variables and the PCA, which showed that species with high levels of damage following drought treatment were the least damaged by the frost treatments (Fig. 1). We assessed frost tolerance using the whole-plant responses survival and regrowth. However, because all species grew well following drought treatment, we were not able to use similar whole-plant responses for drought tolerance. Instead, we used electrolyte leakage, as previous studies have shown that this is a good proxy for drought tolerance measured as survival and regrowth (Bajji, Kinet and Lutts (2002). Therefore, the different measures of drought and frost responses are comparable. Furthermore, we found negative correlations between electrolyte leakage (conductivity) and photosynthetic capacity following drought treatment and between electrolyte leakage and regrowth following frost exposure at both −1 °C and –3 °C. The PCA plots also show covariation among different measured responses to frost and drought treatments, respectively (Fig. 1). Thus, there are expected and reliable signals in our experimental data, meaning that the negative correlation between frost and drought tolerance found is unlikely to be an artefact of the experimental variables used. Instead these results reflect different adaptations to different environments in different species.

### Is leaf dry matter the link between frost and drought tolerance?

We found a significant negative relationship between leaf dry matter content and electrolyte leakage in response to drought, indicating that the former may confer drought tolerance in Pooideae. These results are in line with Liu *et al.,* (Liu *et al*., 2015), who found leaf dry matter content to be positively correlated with drought tolerance in *Magnolia*. We did not, however, find any relationship between leaf dry matter content and regrowth following frost treatment. This indicates that leaf dry matter content is not a component of short term frost responses in Pooideae. This contrasts with Watcharamongkol (Watcharamongkol, 2019), who found a correlation between episodic frost tolerance and water content (the inverse of dry matter content) in the predominantly (sub)tropical PACMAD grasses. One explanation for these contrasting results could be that if high dry matter content was an exaptation to frost tolerance, such that it facilitated adaptation to freezing climates in drought tolerant plants, the signature of that exaptation may be masked by the more sophisticated and complex freezing adaptations of present day temperate clades like Pooideae (Schubert *et al*., 2019b). Thus, it is still possible that the first responses to episodic frost in an ancestor of Pooideae utilised early desiccation tolerance traits, including high dry matter content. Further study of the role of leaf dry matter content in the evolution of drought and frost responses at broader phylogenetic scales and tests for shared gene expression patterns linked to early dehydration traits, including dry matter content, is needed.

### Evolutionary trajectories of drought and frost responses

If drought tolerance had acted as a precursor for frost tolerance, we would have expected drought tolerance to have evolved first in lineages tolerant of episodic (short-term/diurnal) frost, and/or frost tolerance to have evolved more frequently in ancestrally drought tolerant lineages. This is not what we found (Fig. 2). Instead, we found a mirrored phylogenetic pattern for drought and frost tolerance, with the lineage with the highest inferred ancestral drought tolerance (Stipeae) being the least frost tolerant and the highest frost tolerance being measured for species with the lowest ancestral drought tolerance (Poeae + Triticeae). This would suggest that the sophisticated drought and frost tolerance responses of extant species have evolved as independent evolutionary trajectories. These results corroborate findings from a comparative analysis of frost and drought tolerance inferred from Köppen-Geiger zones across all grasses (Schat *et al*., unpubl.) and previous work suggesting that present-day grasses tend to be either frost or drought specialists (Visser *et al*., 2014). These findings are also in line with the idea that there is a tradeoff among abiotic stress responses, such that plants cannot be equally well adapted to multiple stressors, in particular both low temperature and drought (Puglielli, Hutchings and Laanisto, 2021). Finally, our results mean that the evolutionary origins of shared genetic responses to cold and drought remain largely unknown. Gene expression patterns suggest some kind of shared ancestral response to both cold and drought in Pooideae (Schubert *et al*., 2019a). These may have originated even further back in time, outside the Pooideae. Further research in a phylogenetic framework, with species sampled from across broad clades, will be needed to test this further.

Interestingly, we inferred the ancestor of Pooideae to have low frost tolerance, with higher levels being reconstructed only in core Pooideae (Triticeae and Poeae; Fig. 2). This contrasts with other reconstructions, suggesting frost tolerance at deeper nodes, e.g. as far back as the ancestor of all Pooideae except *Brachyelytrum* and *Nardus* plus *Lygeum* (Schubert *et al*., 2019b)(Schat et al. unpubl). The use of an experimentally measured frost response here has thus provided a more nuanced view of how frost tolerance evolved in Pooideae, compared to studies relying on binary coding of this trait (frost sensitive/tolerant). Our reconstruction suggests frost tolerance increased only after transitions to open habitats occurred (Elliott *et al*., 2023; Zhang *et al*., 2022) and coincidentally with the novel expansion of gene families involved in low temperature stress responses in core Pooideae (Sandve and Fjellheim, 2010; Schubert *et al*., 2019a).

We found higher phylogenetic signal to frost than drought tolerance (Table 2), i.e., a higher signature of shared ancestry in frost tolerance than drought tolerance. This holds true even when the effects of phylogeny, geography and the local climate are modelled together (Table 4), meaning it is not an artefact of not accounting for other potentially confounding variables (Freckleton and Jetz, 2009) (Coelho *et al*., 2019; Humphreys and Linder, 2013; Lancaster and Humphreys, 2020). The high phylogenetic signal in frost tolerance is consistent with other studies in grasses (Edwards and Smith, 2010; Humphreys and Linder, 2013)(Schat et al. unpubl.) and land plants (Lancaster and Humphreys, 2020) and suggests that rare evolutionary events structure frost responses in plants.

The low phylogenetic and geographical signals in drought responses (Tables 2 and 4) could indicate that Pooideae rely on general stress tolerance mechanisms to cope with drought, rather than being drought specialists. Alternatively, the low phylogenetic signal suggests that drought tolerance is evolutionarily labile in Pooideae. This is supported by other similarly shallow reconstructions of drought tolerance in Pooideae, with xerophytes arising multiple times independently, only being reconstructed ancestrally in Triticeae (Zhang *et al*., 2022), and drought tolerance being reconstructed at deeper ancestral nodes only in Stipeae and Triticeae (Schat et al. unpubl.). Our results together with these previous reconstructions suggest that origins of modern-day drought tolerance are decoupled from the transitions from closed to open habitats in Pooideae.

Evolutionary lability of drought tolerance is supported by other lines of evidence. Some form of drought tolerance is assumed to have been a key factor in the early evolution of land plants (Oliver, Tuba and Mishler, 2000; Zhao *et al*., 2019), but since then various forms of adaptations for avoiding dehydration have evolved, been lost and regained several times (Bowles, Paps and Bechtold, 2021). Some gene families (e.g. C-REPEAT BINDING FACTORS (CBFs) and dehydrins) are expressed in response to a range of stressors, including drought, frost and salinity (Chew and Halliday, 2011). These gene families are present in all plants at all times and are often larger and more flexible (undergoing expansions and contractions) than gene families not involved in stress responses (Panchy, Lehti-Shiu and Shiu, 2016; Schubert *et al*., 2019a) (Chew and Halliday, 2011). Such flexibility serves as a basis for adaptation, allowing individual genes to be co-opted into different stress responses in certain lineages (Schubert *et al*., 2019a). If this is the genetic basis of drought tolerance in Pooideae, then this evolutionary lability will limit our ability to reconstruct the order in which drought tolerance traits evolved relative to other stress responses and assess their putative roles as evolutionary precursors (Bromham, 2014).

### Local climate conditions do not explain measured drought and frost responses

We found at best a weak correlation between experimentally measured drought and frost tolerance and aspects of the local climate in species’ native ranges (Tables 3, S2). There was no effect of the local climate for frost tolerance or leaf dry matter content but for drought tolerance there was a weak effect of mean annual temperature and minimum temperature of the coldest month. In other words, these results indicate that the most drought tolerant species are from areas that are warm on average but with cold winters. Weak trait-environment relationships are consistent with previous findings for a range of plant traits (upper and lower thermal tolerances, plant height, leaf size, seed size; (Moles *et al*., 2014); (but see (Das *et al*., 2021; Lancaster and Humphreys, 2020)). In ours and previous studies, local temperature conditions explained more of the variation in functional or experimentally determined response variables than local precipitation conditions but neither explained very much. This suggests either that plants in general show only weak signatures of local adaptation and/or that air temperatures expressed by the BioClim variables at coarse geographical scales do not capture the (micro)climatic conditions plants are naturally exposed to (Greiser *et al*., 2020). In support of the latter, land surface temperatures based on satellite measurements of radiative temperatures capture more differences in occupied thermal environments between closely related C3 and C4 grasses than air temperatures (Still, Pau and Edwards, 2014). Another factor not considered here is the stability of trait-environment relationships over time (Cui, 2024; Famiglietti *et al*., 2023). Since our study concerns change in plant traits over evolutionary timescales, incorporating climate fluctuations through time should strengthen trait-environment relationships.

## Conclusion

We conclude that there is little evidence in our data for a positive correlation between drought and frost responses or that drought tolerance acted as a precursor to frost tolerance. Instead, our reconstructions suggest that present-day drought and frost responses are the result of independent evolutionary trajectories in different Pooideae lineages, or that their shared origins occurred outside the Pooideae. Either way, the evolutionary origins of the known physiological and genetic links between frost and drought responses remain unclear. Our results also suggest that origins of modern-day drought tolerance are decoupled from the transitions from closed to open habitats in Pooideae – but also that drought tolerance is more evolutionarily labile than frost tolerance. This lability could limit our ability to reconstruct the relative order in which drought and frost responses originated, potentially hampering assessment of their putative roles as evolutionary precursors. Further research is needed to discern whether our findings are unique to the cool season grasses, or whether signatures of shared evolutionary origins among diverse abiotic stress tolerance responses are no longer detectable on the long timescales studied here.

## Supplementary data

**Supplementary table 1.** Summary of number of plants in experiments.

**Supplementary data 1.** The data input for analyses.

**Supplementary data 2.** Raw data from the experiments.

## Acknowledgements

We are grateful to Mika Kirkhus, Martine Molland, Ane Charlotte Hjertaas, Martin Paliocha, Darshan Young and Øyvind Jørgensen for help with the growth experiment and to two anonymous reviewers for comments on an earlier version of the manuscript.

## Author contributions

SF, JCP and AMH conceived and designed the study; SPS, RW and CLL performed the experiments; SPS, RW, LS and AMH compiled and analyzed the data; all authors interpreted the data; SPS, AMH and SF wrote the paper with input from all authors.

## Data availability

The input for analyses can be found in Supplementary data 1 and the raw data from the experiments can be found in Supplementary data 2.

